# PrediXcan: Trait Mapping Using Human Transcriptome Regulation

**DOI:** 10.1101/020164

**Authors:** Eric R. Gamazon, Heather E. Wheeler, Kaanan P. Shah, Sahar V. Mozaffari, Keston Aquino-Michaels, Robert J. Carroll, Anne E. Eyler, Joshua C. Denny, GTEx Consortium, Dan L. Nicolae, Nancy J. Cox, Hae Kyung Im

## Abstract

Genome-wide association studies (GWAS) have identified thousands of variants robustly associated with complex traits. However, the biological mechanisms underlying these associations are, in general, not well understood. We propose a gene-based association method called PrediXcan that directly tests the molecular mechanisms through which genetic variation affects phenotype. The approach estimates the component of gene expression determined by an individual’s genetic profile and correlates the “imputed” gene expression with the phenotype under investigation to identify genes involved in the etiology of the phenotype. The genetically regulated gene expression is estimated using whole-genome tissue-dependent prediction models trained with reference transcriptome datasets. PrediXcan enjoys the benefits of gene-based approaches such as reduced multiple testing burden, more comprehensive annotation of gene function compared to that derived from single variants, and a principled approach to the design of follow-up experiments while also integrating knowledge of regulatory function. Since no actual expression data are used in the analysis of GWAS data - only *in silico* expression - reverse causality problems are largely avoided. PrediXcan harnesses reference transcriptome data for disease mapping studies. Our results demonstrate that PrediXcan can detect known and novel genes associated with disease traits and provide insights into the mechanism of these associations.

## INTRODUCTION

Genome-wide association studies (GWAS) have been remarkably successful in identifying susceptibility loci for complex diseases. These studies typically conduct single-variant tests of association to interrogate the genome in an agnostic fashion and, due to modest effect sizes, have come to rely on ever-greater sample sizes^1,2^ to make meaningful inferences. We have been less successful in developing methods that improve on existing simple approaches. In general, the genetic associations identified as genome-wide significant thus far account for only a modest proportion of variance in disease risk^3^. Indeed, there is now widespread recognition, if not consensus, that GWAS of disease susceptibility (for which, the relevant genetic effects may be very small) and pharmacologic traits (for which large effect sizes are not unusual)^4,5^ have resulted in limited conclusive findings on the genetic factors contributing to complex traits. Importantly, the functional significance of most discovered loci, including even those that have been the most reproducibly associated, remains unclear. Assigning a causal link to the nearest gene falls short of elucidating a functional connection, as recently demonstrated by the obesity-associated variants within *FTO* that form longrange functional connections with *IRX3*^6^. And while GWAS will no doubt continue to identify many more susceptibility loci, the question of how to advance biological knowledge of the underlying mechanisms of disease risk remains a paramount challenge.

A large portion of phenotypic variability in disease risk for a broad spectrum of disease phenotypes can be explained by regulatory variants, i.e. genetic variants that regulate the expression levels of genes^7-10^. For example, almost 80% of the chip-based heritability of disease risk for 11 diseases from the WTCCC can be explained by genome variation in DNase I hypersensitivity sites, which are likely to regulate chromatin accessibility and thus transcription^11^.

Large genomic consortia (e.g., ENCODE^12^) are generating an unprecedented volume of data on the function of genetic variation. The Genotype-Tissue Expression (GTEx^13^) project is an NIH Common Fund project that aims to collect a comprehensive set of tissues from 900 deceased donors (for a total of about 20,000 samples) and to provide the scientific community a database of genetic associations with molecular traits such as mRNA levels. (See GTEx main paper^14^ on Phase 1 data.) Other large-scale transcriptome datasets include Genetic European Variation in Health and Disease^15^ (GEUVADIS, 460 lymphoblastoid cell lines), Depression Genes and Networks (DGN, 922 whole blood samples)^16^, and Braineac (130 individuals with multiple brain region samples)^17^. Yet, effective methods that harness these reference transcriptome datasets for disease mapping are lacking.

Methodologically, gene-based approaches and multi-marker association tests have been developed as alternatives to traditional single-variant tests. By conducting tests of association on biologically informed aggregates of SNPs, such tests seek to evaluate *a priori* functionally relevant units of the genome and, in many cases, reduce the multiple-testing penalty that plague single-variant approaches, by 10 to 100 fold. The incorporation of -omics data, such as those being generated by high-resolution transcriptome studies, provides a means to extend genome-wide association studies by addressing the functional gap. Technological advances in high-throughput methods have reinforced the important finding that intermediate molecular phenotypes are under significant genetic regulation, with expression quantitative trait loci (eQTLs) as the predominant example. However, approaches that fully leverage the comprehensive regulatory knowledge generated by transcriptome studies are relatively lacking despite the fact that these studies have the potential to dramatically improve our understanding of the genetic basis of complex traits^13^.

We hypothesized that a SNP aggregation approach that integrates information on whether a SNP regulates the expression of a gene can greatly increase the power to identify trait-associated loci either from a strong functional SNP signal or from a combination of modest signals, the so-called grey area of GWAS. The present study suggests that PrediXcan, a novel method that incorporates information on gene regulation from a set of markers, increases the power to detect associations relative to traditional SNP-based GWAS and known gene-based tests under a broad range of genetic architectures and provides mechanistic insights and more easily interpreted direction of effect into the observed associations.

## RESULTS

### PrediXcan method

PrediXcan, by design, exploits genetic control of phenotype through the mechanism of gene regulation as a way to identify trait-associated genes. Figure 1 is a schematic diagram of the regulatory mechanism that is tested with PrediXcan. An individual’s gene expression level (typically unobserved in a GWAS) is decomposed into a genetically regulated expression (*GReX*) component, a component altered by the trait itself (i.e., a reverse causal effect that may occur if disease status or other conditions alter expression levels), and the remaining component attributable to environmental and other factors. PrediXcan tests the mediating effect of gene expression by quantifying the association between *GReX* and the phenotype of interest.

**Figure 1.**
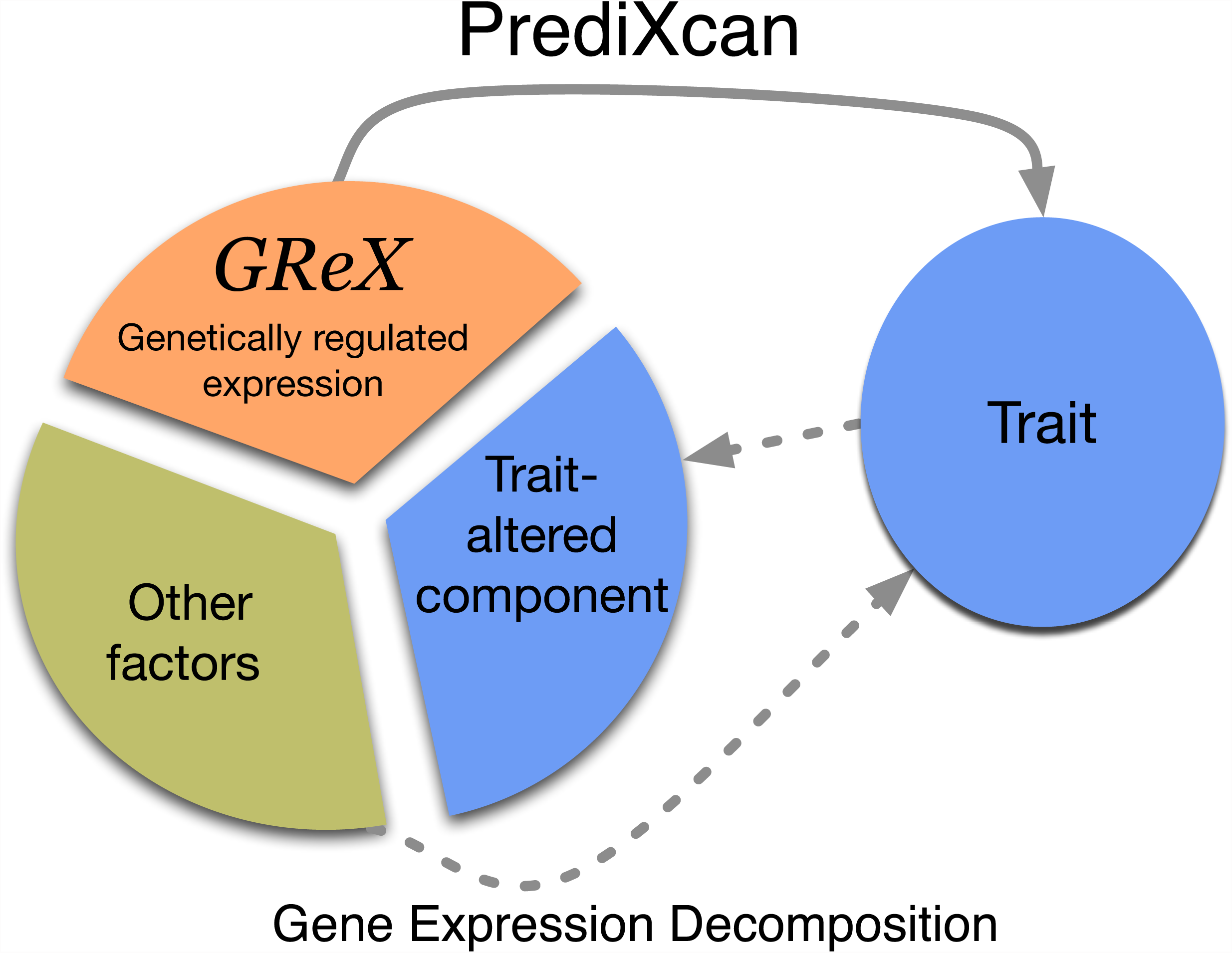
Mechanism tested by the PrediXcan method. This figure shows the conceptual decomposition of the expression level of a gene into three components: genetically determined component, a component altered by the trait itself, and the remaining factors (including environment). PrediXcan estimates the genetically regulated component of expression *(GReX*) and correlates it with the trait to identify trait-associated genes.

We use reference transcriptome datasets from studies such as GTEx^13^, GEUVADIS^15^, and DGN^16^ among others, to train additive models of gene expression levels. These models allow us to estimate the genetically regulated expression, *GReX*. We denote the estimated value with a hat, 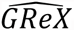 These estimates constitute multiple-SNP prediction of expression levels. The weights for the estimation are stored in our publicly available database.

The analogy with genotype imputation is relevant here. Genotype imputation uses information from a reference sample to learn how to impute genotypes at the unmeasured SNPs in the test set. Similarly, PrediXcan uses a reference dataset in which both genome variation and gene expression levels have been measured to develop prediction models for gene expression. We use these prediction models to “impute” gene expression (which is unobserved in a typical GWAS), and we do so by estimating the genetically determined component, *GReX*.

PrediXcan application to a GWAS dataset consists of “imputing” the transcriptome using the weights derived from reference transcriptome datasets and correlating the *GReX* with the phenotype of interest using regression methods (e.g., linear, logistic, Cox) or non-parametric approaches (e.g., Spearman). (For the specific results on disease phenotypes analyzed here, we used logistic regression with disease status.) We are aware of the attenuation bias that arises because of the error in the estimation of *GReX.* This is a subject to be investigated in the future, but this bias does not invalidate our analysis since we only use the estimate of *GReX* as a discovery tool. Figure 2 summarizes the flow of the method development described above.

**Figure 2.**
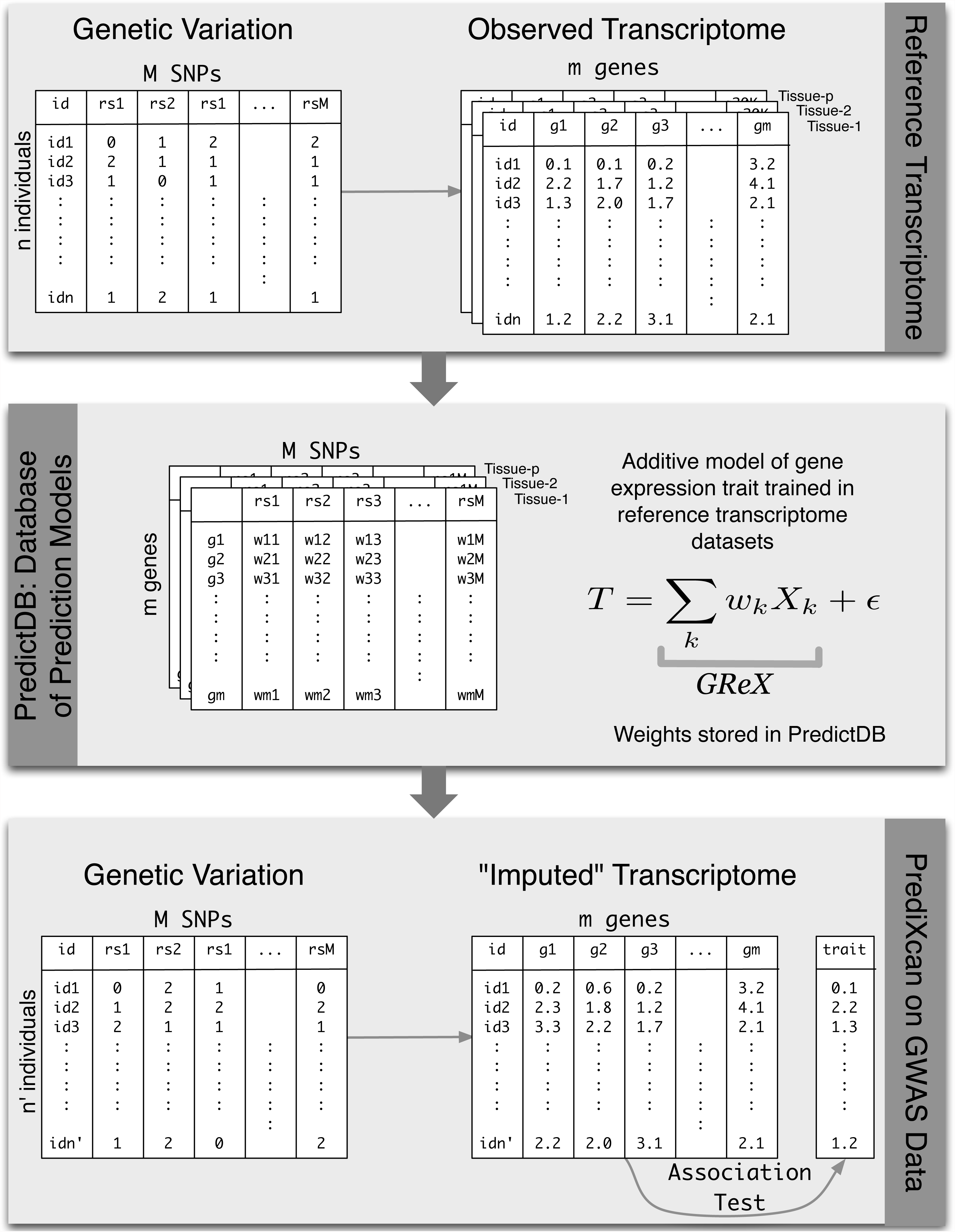
PrediXcan framework. The workflow illustrates the steps used in developing the PrediXcan method. The top panel shows the data used from the reference transcriptome studies: genotype and expression levels (GTEx, GEUVADIS, DGN, etc). The sample size of the study is denoted by *n*, *m* is the number of genes considered, *M* is the total number of SNPs, and *p* is the number of available tissues. The second panel shows the additive model used to build a database of prediction models, PredictDB. *T* represents the expression trait, and *X*_*k*_ is the number of reference alleles for SNP *k*. The coefficients of the models for each tissue are fitted using the reference transcriptome datasets and optimal statistical learning methods chosen among LASSO, Elastic Net, OmicKriging, etc. The bottom panel shows the application of PrediXcan to a GWAS dataset. Using genetic variation data from the GWAS and weights in PredictDB, we “impute” expression levels for the whole transcriptome. These imputed levels are correlated with the trait using regression (e.g., linear, logistic, Cox) or non-parametric (Spearman) approaches. (For the disease phenotypes in the WTCCC datasets and the replication dataset reported here, we used logistic regression with disease status.)

### Features of PrediXcan

PrediXcan is, as we have emphasized, particularly focused on a mechanism – gene expression regulation – that has already been established as being contributory to common diseases, including psychiatric and neurodevelopmental disorders^7^. The test has the potential to identify gene targets for therapeutic applications because it is inherently mechanism-based and provides directionality.

Additional advantages include:

- Like other gene-based tests, it has much smaller multiple-testing burden (∼20K tests maximum, ∼10K genes with high quality prediction in most tissues) compared to single variant tests (∼5-10M tests). Moving beyond the stringent Bonferroni correction, priors on genes can be less restrictive than for SNPs.
- Informative priors and groupings of functional units (based on known pathways, for example) are much more straightforward to construct for genes than SNPs.
- No actual transcriptome data are required since the predicted expression levels are a function of genetic variation alone. Thus, the method can be applied to any existing dataset with large-scale genome interrogation such as those in dbGaP or other repositories. Re-analyses of existing datasets, with a focus on mechanism using PrediXcan, address a gap that has largely characterized GWAS to date.
- Reverse causality is not a major concern since disease status or drug treatment does not alter germline genomic variation.
- Meta-analysis of gene-based results is simplified since less stringent harmonization between studies is required.
- Multiple tissues can be evaluated using a reference transcriptome dataset (such as GTEx). In general, the only limitation is the availability of gene expression data in the given tissue for model building, which need not be, from the same study as that used for phenotype investigation. In cases where transcriptome data are available, separate analyses should be performed to simplify interpretation.
- The approach can be applied to common or rare variants. In general, larger sample sizes for the training set will be needed to achieve good prediction models with rare variants.

### Publicly available database of prediction models and software

We make the prediction models (derived from LASSO^18^ and elastic net^19^) and the software to predict the transcriptome (in a variety of tissues) (see Materials and Methods) publicly available (see https://github.com/hakyimlab/PrediXcan).

### Predicting the transcriptome

#### Prediction model selection

We built prediction models in the DGN whole blood cohort using LASSO, the elastic net (*α*=0.5), and polygenic score at several p-value thresholds (single top SNP, 1e-04, 0.001, 0.01, 0.05, 0.5, 1). We assessed predictive performance using 10-fold cross-validation (*R*^*2*^ of estimated *GReX* vs. observed expression) as well as in an independent set. We found that LASSO performed similarly to the elastic net and that LASSO outperformed the polygenic score at all thresholds, although all methods are highly correlated (see Supplemental Figure 1). For subsequent analyses, we focused on the prediction models using the elastic net because we found it to perform well and to be more robust to slight changes in input SNPs (potentially due to variations in imputation quality between cohorts).

#### Heritability estimation and comparison with prediction R^2^

We estimated the heritability of gene expression in DGN attributable to SNPs in the vicinity of each gene using a mixed-effects model (see Materials and Methods) and calculated variances using restricted maximum likelihood as implemented in GCTA^20^. We use only local SNPs since we found that heritability estimates using all genotyped SNPs were too noisy to make meaningful inferences.

We use heritability estimates as our benchmark for the prediction *R*^*2*^ since this constitutes the upper limit of our prediction performance. For genes for which an elastic net model was available (n=10,427), the average heritability in DGN was 0.153. In comparison, the average 10-fold cross-validated prediction *R*^*2*^ for elastic net was quite close at 0.137; for the polygenic score (P<1 × 10^−4^) and top-SNP models, average prediction *R*^*2*^ values were sizably lower at 0.099 and 0.114, respectively. We show the performance *R*^*2*^ for each model in Figure 3, with the corresponding heritability estimate and confidence interval in the background for comparison. We also note that elastic net predictive performance reached or exceeded the lower bound of the heritability estimate for 94% of genes, while polygenic score (P<1 × 10^−4^) did so for just 76% of the genes and the top SNP for 80% of the genes (Figure 3), consistent with the performance ranking given by the average (across genes) *R*^*2*^.

**Figure 3.**
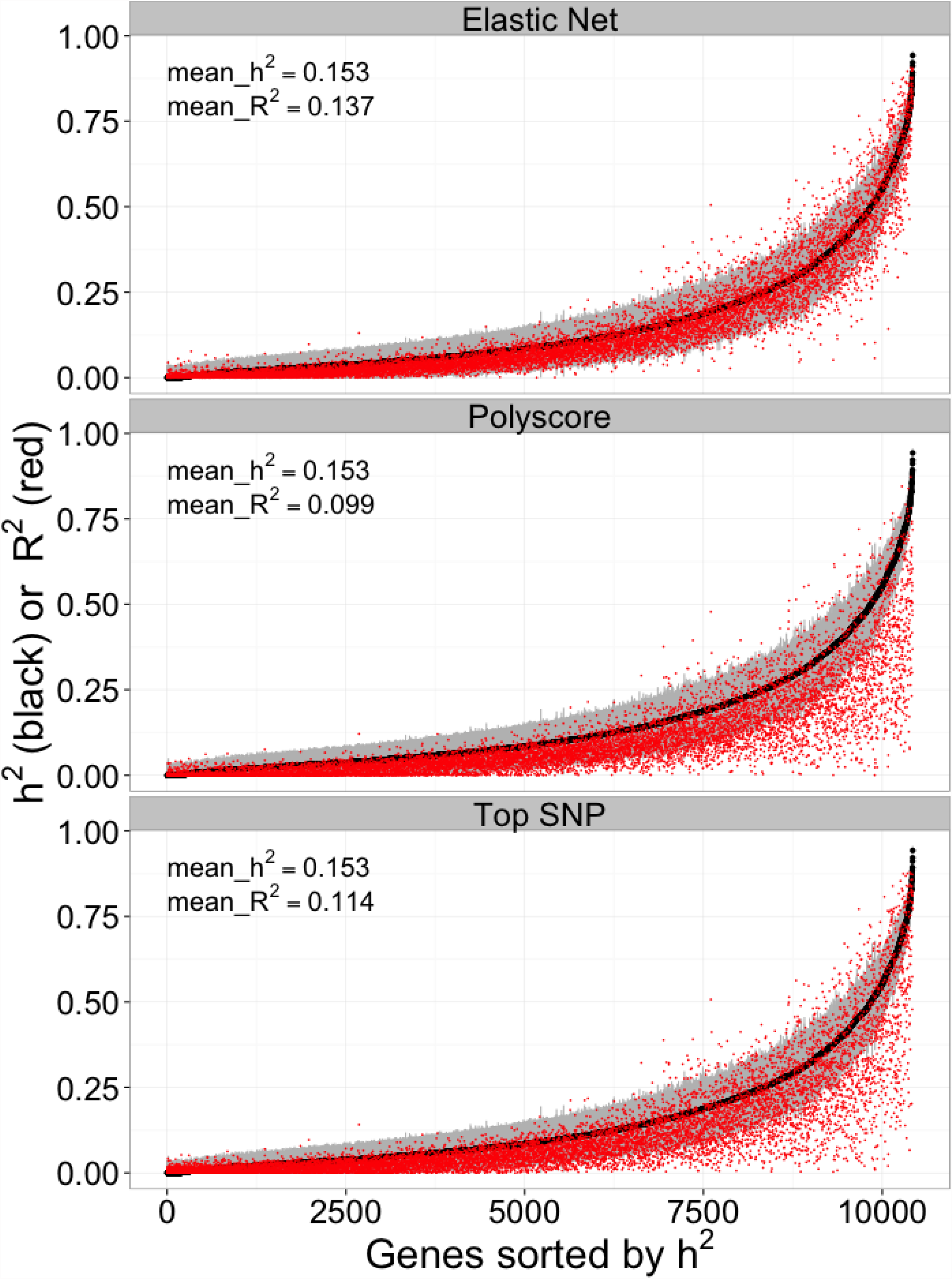
Cross-validated prediction performance vs heritability. This figure shows the prediction performance (R2 of GReX vs. observed expression in red) compared to gene expression heritability estimates (black with 95% confidence interval in gray). Performance was assessed using 10-fold cross-validation in the DGN whole blood cohort (n=922) with the elastic net, polygenic score (p < 1e-04), and using the top SNP for prediction.

#### Comparing imputed vs. directly genotyped SNPs as reference set for prediction model

Predictive performance of elastic net was similar whether all SNPs from the 1000 Genomes imputation or the HapMap Phase II subset were included in the model building (Supplemental Figure 2). Models based on imputed data (both the 1000 Genomes and the HapMap subset) substantially outperformed models based on genotyped SNPs in WTCCC (Supplemental Figure 2). Thus, we chose the elastic net models built in the smaller HapMap SNP subset, relative to 1000 Genomes, in our applications of PrediXcan to reduce computation time without sacrificing performance. As reference transcriptome studies increase in sample size, we may need to switch to a more dense imputation to take advantage of increased prediction performance from rare variants.

#### Prediction performance in an independent test set

We also tested the prediction models trained in the DGN whole blood cohort on several independent test cohorts with available whole-genome genotype and transcriptome data. We used weights derived from the DGN whole blood data (“training set”) to predict gene expression levels (treated as quantitative traits) in GEUVADIS LCLs (lymphoblastoid cell lines) and nine GTEx pilot tissues (“test sets”). Figure 4 provides a Q-Q plot showing the expected (under the null, correlation between two independent vectors with the same sample size) and observed *R*^*2*^ (between observed and predicted) from the elastic net prediction in GEUVADIS LCLs. We find a substantial departure from the null distribution indicating that the elastic net model trained in DGN (equation 2 of Materials and Methods, with effect size estimates 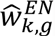) captures a significant proportion of the transcriptome variability. The average prediction *R*^*2*^ is 0.0197 for GEUVADIS LCLs. For GTEx tissues, the prediction *R*^*2*^ values are 0.0367 (adipose), 0.0358 (tibial artery), 0.0356 (left ventricular heart), 0.0359 (lung), 0.0269 (muscle), 0.0422 (tibial nerve), 0.0374 (sun exposed skin), 0.0398 (thyroid), and 0.0458 (whole blood). Interestingly, we also find a substantial departure from the null distribution of expected *R*^*2*^ values for predicted expression using DGN weights in each of the nine GTEx tissues suggesting that models developed in whole blood are still useful for understanding diseases that affect other primary tissues (Supplemental Figure 3). Consistent with this, average prediction *R*^*2*^ is highest for whole blood as expected but the loss in power for other tissues is modest.

**Figure 4.**
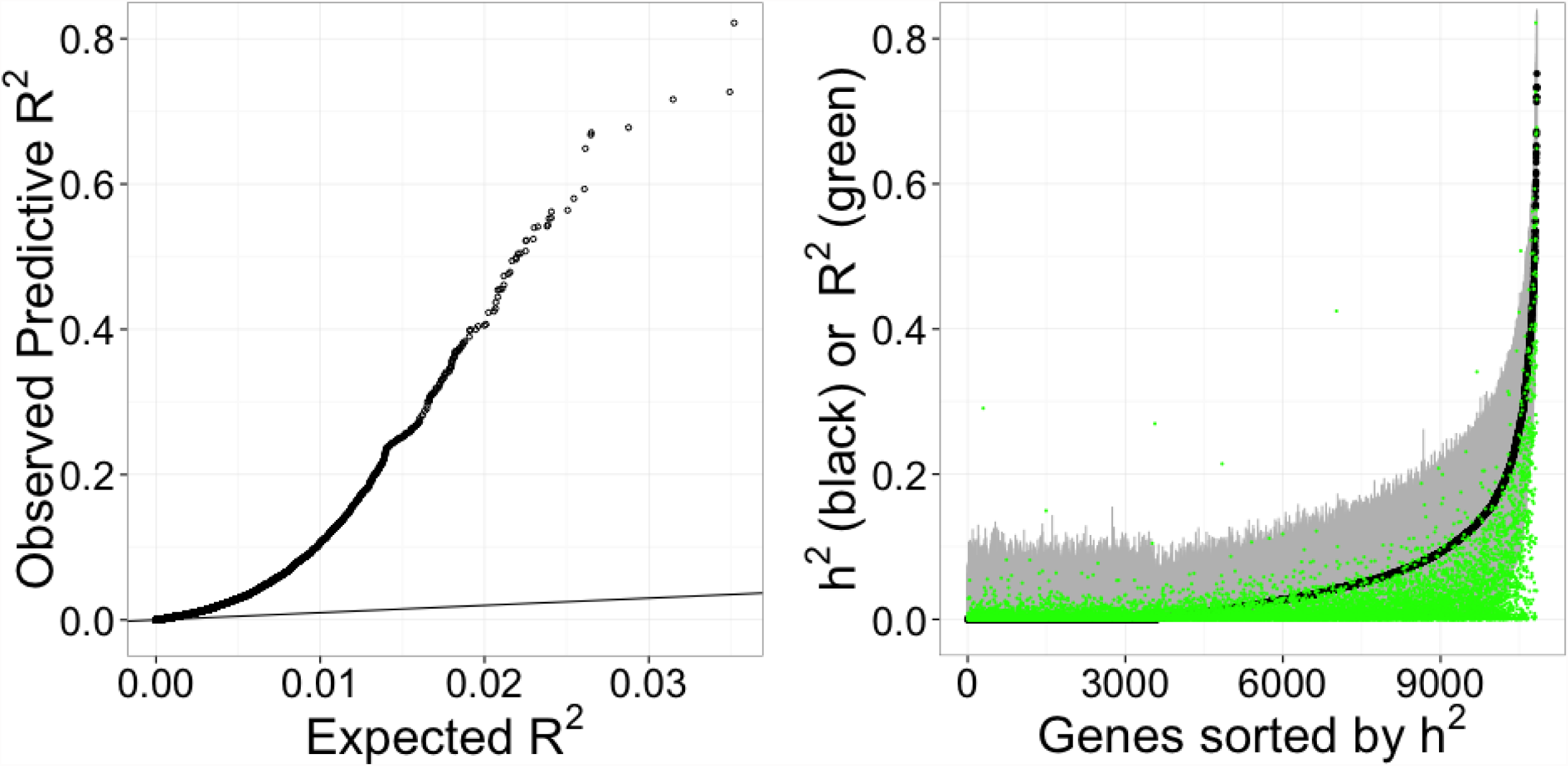
Prediction performance of elastic net tested on a separate cohort. Using whole blood prediction models trained in DGN, we compared predicted levels of expression with observed levels on lymphoblastoid cell lines from the 1000 Genomes project. RNA-sequenced data (n=421) on these cell lines have been made publicly available by the GEUVADIS consortium. Left panel shows the squared correlation, *R2*, between predicted and observed levels plotted against the null distribution of *R2*. Right panel shows prediction performance (R^2^ of GReX vs. observed expression in green) compared to GEUVADIS gene expression heritability (h^2^) estimates (black with 95% confidence interval in gray).

#### Examples of well-performing genes

Figure 5 illustrates the genes with some of the highest correlations from this analysis, providing a comparison of the predicted expression and the observed expression. Among these genes, both *ERAP2* and its paralog *ERAP1* play fundamental roles in MHC antigen presentation^21^, immune activation and inflammation.

**Figure 5.**
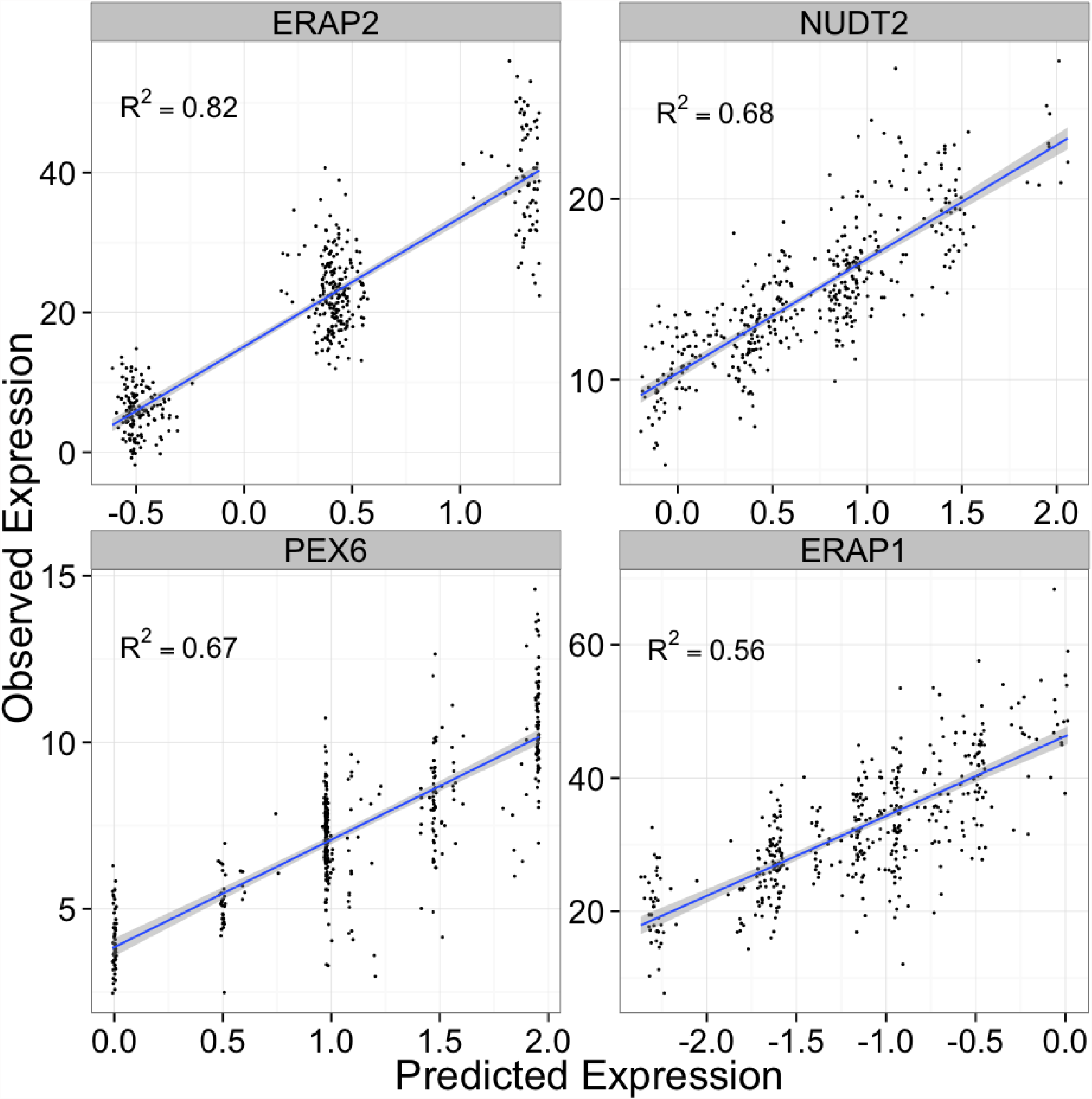
Examples of well-predicted genes. These plots show observed vs. predicted levels of 4 genes. Predicted levels were computed using whole blood elastic net prediction models trained in DGN data. Observed levels were RNA-seq data in lymphoblastoid cell lines generated by the GEUVADIS consortium.

#### Marginal gain in performance when adding distant SNPs

We also generated prediction models trained in the DGN whole blood cohort that included *trans*-eQTLs (>1Mb from gene start or end or on a different chromosome) generated from linear regression (p<10^−5^). We tested the predictive performance of these models in the GTEx whole blood cohort. While a few genes had higher correlations between predicted and observed expression than expected by chance, the departure from the null distribution was much smaller than that for the prediction models based on local SNPs (Supplemental Figure 4), perhaps due to the low power to map *trans* SNPs. Based on this result, in this paper we focus primarily on results based on local SNPs.

### Application of PrediXcan to WTCCC

We applied PrediXcan to seven complex disease phenotypes from the WTCCC study^22^. For this purpose, we utilized the DGN whole blood elastic net prediction models. We correlated the estimated genetically regulated gene expression for close to 8700 genes with disease status for each WTCCC dataset and identified 41 significant associations (Bonferroni corrected *p* < 0.05) with five diseases (Table 1). Notably, we identified 29 genes associated with type 1 diabetes (T1D) risk (Table 1 and Fig. 6), 8 of which were outside of the extended MHC. Complete results for the remaining 6 diseases are shown in Supplemental Figures 5 and 6. Consistent with the original GWAS of WTCCC diseases, our most significant results were for autoimmune diseases^22^.

**Table 1.**
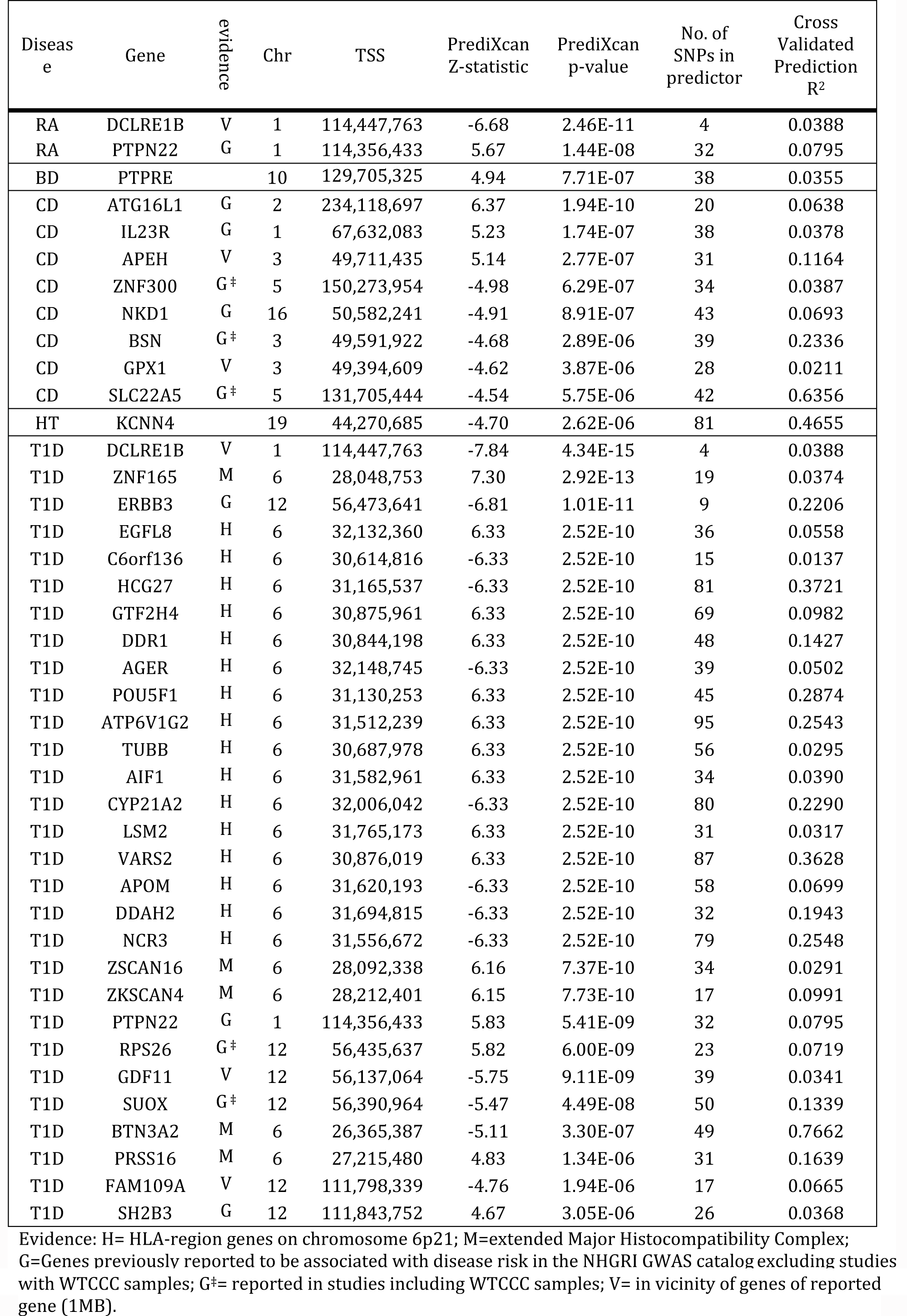
Top PrediXcan results for WTCCC using DGN whole blood prediction models. PrediXcan results for bonferroni significant gene associations. To account for multiple testing, we used a significance threshold of 5.76x10^−6^ for all diseases. Chromosome and gene start position are based on GENCODE version 12. The cross validated prediction *R*^*2*^ between predicted and observed gene expression is based on 10-fold cross validation within the DGN whole blood sample.

**Figure 6.**
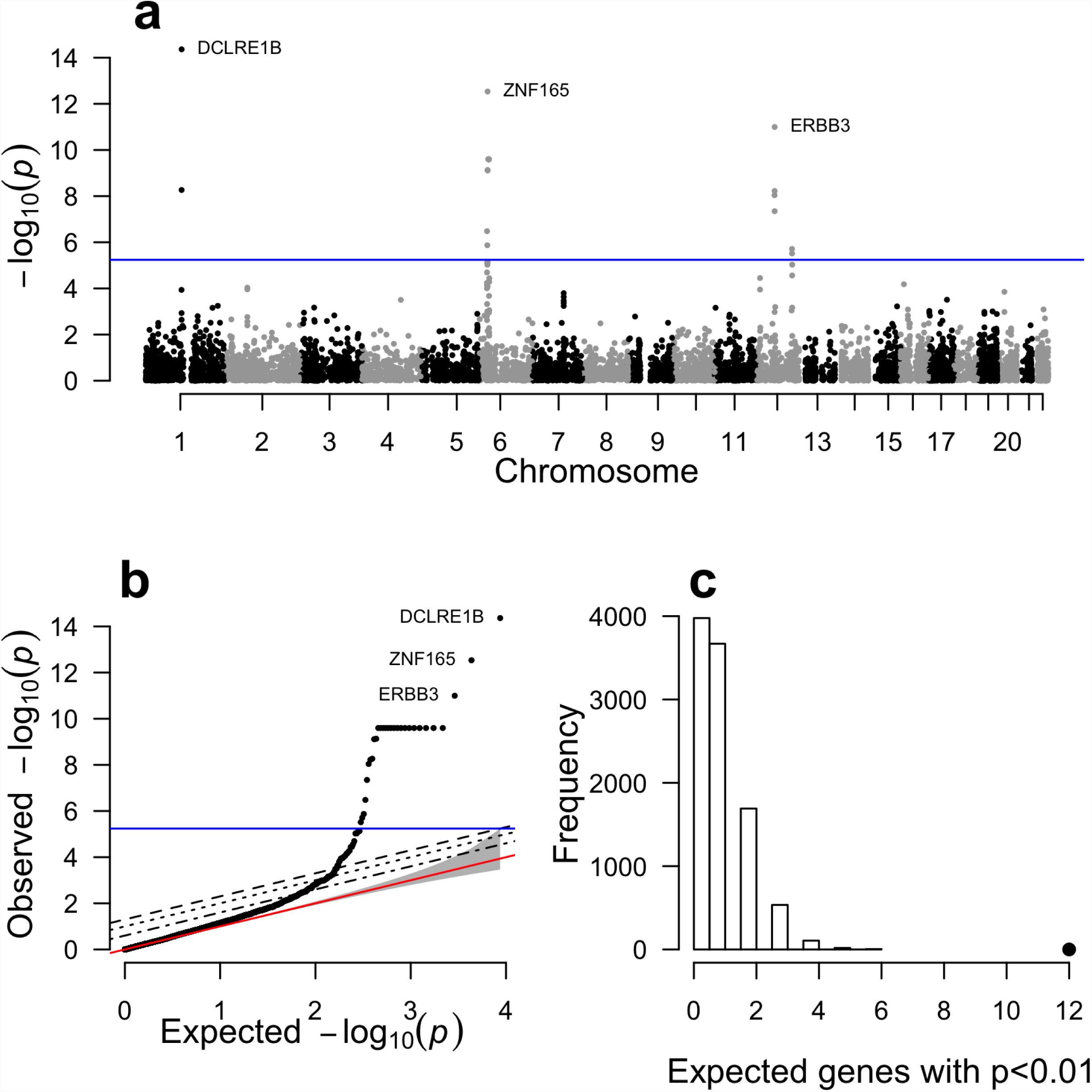
PrediXcan results for type 1 diabetes. Complete results for our analysis of type 1 diabetes from the WTCCC using gene expression predicted with the DGN whole blood predictors. Panel (a) shows association p-values based on gene position across the genome. Panel (b) shows the same results plotted against the null expectation in a q-q plot. The red line in panel (b) shows the null expected distribution of p-values. In panels (a) and (b), the blue line represents the bonferroni corrected genome-wide significance threshold. The top 3 genes are labeled. Panel (c) shows the results of our GWAS enrichment analysis. The histogram shows the expected number of genes with a p-value < 0.01 based on 10,000 random permutations. The large point shows the observed number of previously known T1D genes that fall below this threshold.

#### DCLRE1B and PTPN22

As has been previously reported for complex autoimmune diseases^23^, we observed genes that were associated with multiple autoimmune diseases, namely T1D, Crohn’s disease (CD), and rheumatoid arthritis (RA). Interestingly, the top (genome-wide significant) PrediXcan gene for both T1D and RA, *DCLRE1B*, has not been previously implicated in either disease, but has been linked to CD, ulcerative colitis and inflammatory bowel disease^24^. Lower predicted expression of *DCLRE1B* was associated with increased disease risk for both RA and T1D. Interestingly, higher predicted expression of *DCLRE1B* was nominally associated with increased Crohn’s disease risk in our PrediXcan analysis (p = 0.001). Similarly, *PTPN22* was significantly (positively) associated with RA and T1D (table 1), and nominally (negatively) associated with CD (p-value = 0.017). Previous single variant analyses implicated *PTPN22* with multiple autoimmune diseases including RA, T1D, CD, myasthenia gravis, and vitiligo according to the NHGRI catalog^25^. These results highlight the known overlap in genetic risk factors for autoimmune diseases.

#### Prior evidence of significant genes

All genes in table 1, excluding *PTPRE* and *KCNN4* (discussed below), have been either previously reported with GWAS studies, or are located in the vicinity of reported genes (within 1MB). For T1D, 5 out of the 29 genome-wide significant genes have been reported via conventional single variant analyses (as curated by the NHGRI^25^ repository of GWAS results). Furthermore, 21 of the genes associated with T1D in our analysis lie within the extended MHC (Table 1), a region that is known to be associated with disease risk^26^. Additionally, *ERBB3*, which contains SNPs previously associated with T1D in GWAS^27^, showed a negative correlation with disease risk in PrediXcan (p < 10^−^11), which is consistent with a prior study that showed risk genotypes associated with lower expression of *ERBB3* in PBMCs^28^. Furthermore, it has been reported that subjects with protective genotypes had higher percentages of ERBB3^+^ monocytes and dendritic cells leading to greater T cell proliferation^28^. These results highlight one of the key advantages of PrediXcan, which is to provide the direction of effect.

#### Enrichment of previously reported disease genes

The results described above highlight gene associations that attain genome-wide significance. Additionally, we tested for enrichment of reported disease genes among our PrediXcan results using less stringent significance thresholds. Reported genes were derived from the comprehensive NHGRI catalog of disease-associated variants identified using GWAS^25^. Five of the seven diseases (bipolar disorder (BD), coronary artery disease (CAD), CD, RA, T1D) had a significant enrichment of reported genes in the PrediXcan results (Figure 6C, Supplemental Figure 7). Results for other p-value thresholds were similar (results not shown). These enrichment analyses on the PrediXcan findings suggest that among the genes that fail to meet strict genome-wide significance, there are likely to be true disease associations.

#### Potentially novel findings

In addition to the results described above for autoimmune diseases, we identified two potentially novel disease-associated genes. Lower predicted expression of *KCNN4* was associated with an increased risk of hypertension (p-value = 2.62 x 10^−6^, Table 1) and high predicted *PTPRE* expression was associated with increased risk of bipolar disorder (p-value = 7.71 × 10^−7^, Table 1). Interestingly, an intronic SNP in *PTPRE* was previously found to associate with response to the stimulant amphetamine^29,30^. In contrast to the original WTCCC single-variant analyses^22^, the PrediXcan analysis for bipolar disorder and hypertension produced genome-wide significant results. Additional studies of these genes are warranted.

#### Comparison to large single variant meta-analyses

Using publically available meta-analysis results, we summarized the single-variant association results for SNPs that are included in the prediction models for the top disease-associated PrediXcan genes. As expected, the genes associated with autoimmune diseases (RA and CD) each contained multiple SNPs that are individually associated with disease risk (Supplemental Table 1). Thus, the identified disease gene associations are consistent with the single-variant meta-analysis results. Interestingly, in many cases, we detect these associations with much smaller sample sizes.

Furthermore, our gene-based results allow for more direct biological interpretation compared to individual SNPs.

The PrediXcan associations for BD and HT have not been observed before using traditional single-variant GWAS. The association between predicted expression of *PTPRE* and BD is further supported by single variant meta-analysis results from the Psychiatric Genetics Consortium (PGC)^31^. Table 2 shows the meta-analysis p-values for each SNP included in the predictor of *PTPRE*. While none of the SNPs is individually genome-wide significant, 10 out of 23 are nominally associated with BD disease risk in the PGC meta-analysis (p<0.05). Follow-up studies of this disease association are necessary, but our analysis in combination with existing results suggests *PTPRE* may be an excellent BD candidate gene. Furthermore, this result highlights the advantage of our gene-based approach that combines information across SNPs, each of which many only contribute nominally to disease risk and therefore remain below the detection limits of single-variant analyses.

**Table 2.** Association between *PTPRE* predictor SNPs and bipolar disorder. The 24 SNPs listed below are included in the DGN whole blood prediction model for *PTPRE*. The association p-values for these SNPs in the WTCCC data and the PGC Bipolar disorder meta-analysis are shown.

| SNP | Gene | Meta Analysis P-value* | WTCCC GWAS P-value |
| --- | --- | --- | --- |
| rs10764743 | PTPRE | 9.89E-05 | 6.947E-08 |
| rs2096285 | PTPRE | 0.001868 | 0.00004311 |
| rs2782877 | PTPRE | 0.002848 | 0.2231 |
| rs4353188 | PTPRE | 0.003062 | 0.00006297 |
| rs4568912 | PTPRE | 0.01409 | 0.009569 |
| rs6911 | PTPRE | 0.01929 | 0.01019 |
| rs3186827 | PTPRE | 0.02187 | 0.009356 |
| rs4237449 | PTPRE | 0.02238 | 0.009648 |
| rs4548544 | PTPRE | 0.02276 | 0.009541 |
| rs1127299 | PTPRE | 0.02422 | 0.009205 |
| rs10764815 | PTPRE | 0.0998 | 0.5328 |
| rs11018116 | PTPRE | 0.1343 | 0.1607 |
| rs1273180 | PTPRE | 0.1941 | 0.08975 |
| rs2799399 | PTPRE | 0.2097 | 0.07739 |
| rs1255136 | PTPRE | 0.4116 | 0.09816 |
| rs10764806 | PTPRE | 0.5628 | 0.4339 |
| rs2387863 | PTPRE | 0.6786 | 0.2429 |
| rs4750963 | PTPRE | 0.7605 | 0.3659 |
| rs7916444 | PTPRE | 0.7856 | 0.5639 |
| rs11016020 | PTPRE | 0.8522 | 0.01473 |
| rs2766034 | PTPRE | 0.9328 | 0.8095 |
| rs11017197 | PTPRE | 0.9746 | 0.8206 |
| rs11017201 | PTPRE | 0.9802 | 0.8237 |

#### Comparison of gene-based tests

We applied PrediXcan and two widely-used gene-based tests (VEGAS and SKAT) to WTCCC. In a Q-Q plot showing all three distributions of p-values, for genes outside of the HLA region, from these gene-based tests (Figure 7), SKAT had improved performance relative to VEGAS, and PrediXcan showed the most extreme departure from the null at the tail end of the distribution.

**Figure 7.**
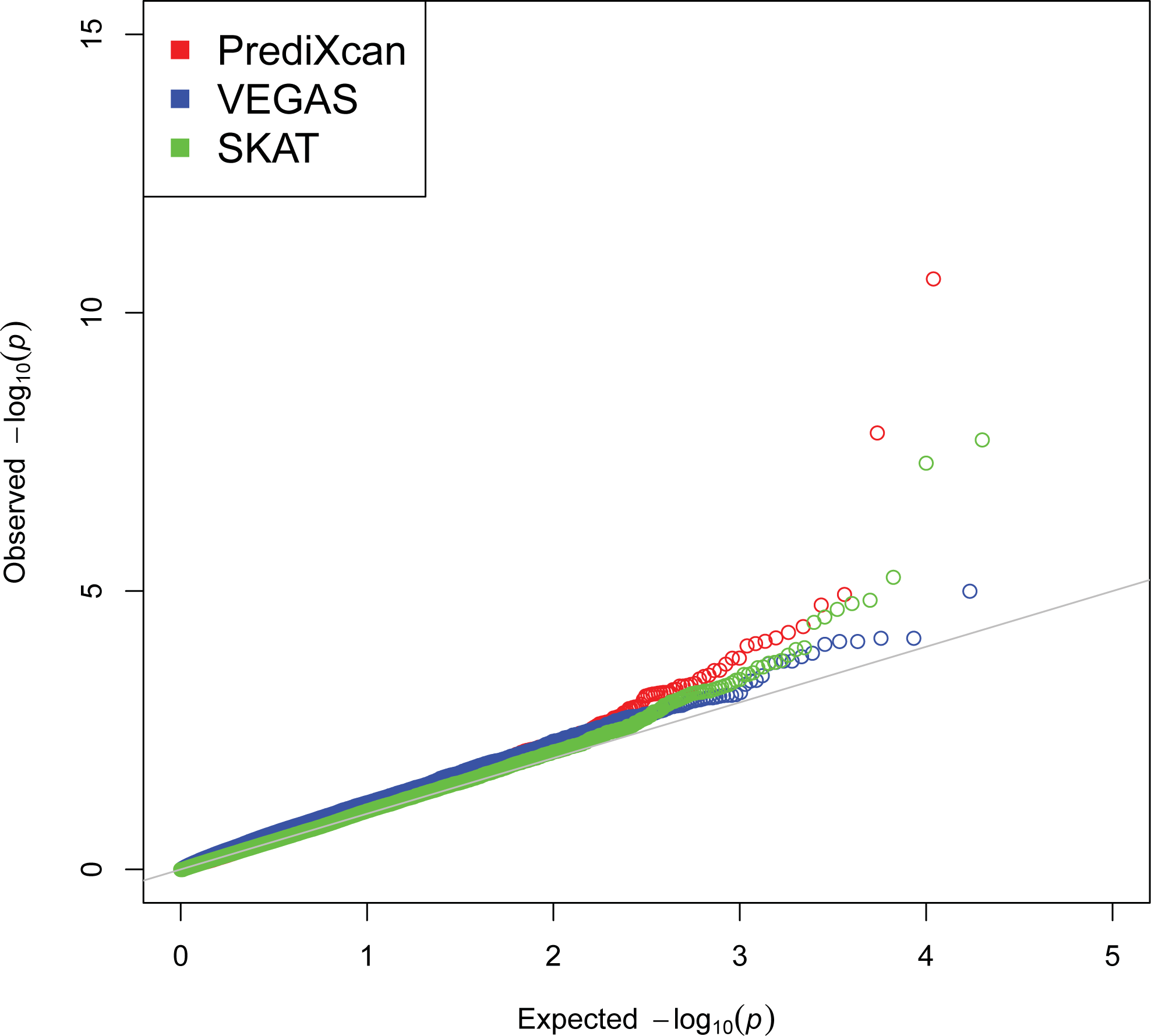
Comparison of gene-based methods. Q-Q plot showing distribution of p-values derived from each method (VEGAS, SKAT, and PrediXcan) for genes outside of the HLA region for Rheumatoid Arthritis.

#### Replication of PrediXcan findings

To replicate our findings, we applied the DGN elastic net whole blood prediction models to an independent rheumatoid arthritis GWAS from Vanderbilt University’s BioVU repository (see Materials and Methods). Both genes (*DCLRE1B* and *PTPN22*) that were found to be genome-wide significant in the WTCCC rheumatoid arthritis data were also significant, with concordant direction of effect, in the replication samples (p = 0.012 and p = 0.036, respectively).

## DISCUSSION

Gene expression, as an intermediate phenotype between genetic variation and higher-level phenotypes, is an important mechanism underlying disease susceptibility and drug response. Studies of the transcriptome in several tissues^13^ have shown that variation in gene expression is heritable^32,33^ and can be mapped to the genome. Particularly, eQTL mapping provides an immediate view of the effects of genetic variants on the phenotype closest to genetic variation, namely transcript abundance, and thus promises to enable the discovery of the molecular mechanisms underlying human phenotypic variation^34^. Furthermore, transcriptome regulation studies facilitate the consideration of thousands of gene expression phenotypes in parallel, thereby enabling a comprehensive approach to understanding the genetic basis of complex traits^35^. In this study, we developed a method that explicitly utilizes the wealth of regulatory information derived from transcriptome regulation studies to map trait-associated loci.

Traditional GWAS test single markers for association with phenotype. This approach ignores a considerable amount of information contained in the genome, including functional information for the markers as well as the structure implied, for example, by the physical contiguity of the loci. The SNPs that are considered in our analyses are likely to regulate the expression of a gene as expression quantitative trait loci, and PrediXcan uses their effect on gene expression to infer the genetically regulated expression level in GWAS samples for which expression data are generally not available. Moreover, the gene-based test we describe here takes into account the potential tissue specificity of the regulatory SNPs.

Our PrediXcan method tests the mediating effects of gene expression levels by quantifying the association between the genetically regulated levels of expression and the phenotype of interest. To implement this, we developed prediction models of gene expression using large-scale transcriptome study datasets (DGN, GEUVADIS, GTEx). After extensive testing, we chose to use the elastic net model, which performed similarly to LASSO, but substantially outperformed simple polygenic approaches. Manor and Segal^36^ have published results on robust prediction of expression levels using K nearest neighbor (KNN) and elastic net approaches. Based on their conclusion that a combination of elastic net and KNN along with the use of genomic annotation such as GC content can improve prediction performance, it is reasonable to hypothesize that the incorporation of a more comprehensive functional annotation approach into the PrediXcan framework can yield additional performance gain.

Application of the method to WTCCC data recapitulated many known loci but also identified novel genome-wide significant genes. We believe that a systematic reanalyses of GWAS datasets in comprehensive repositories such as dbGAP and the European Genome-phenome Archive (EGA) could provide a cost-effective approach to uncovering novel disease mechanisms using only existing genomic resources.

In contrast to other gene-based tests, PrediXcan provides the direction of effect, which may yield opportunities for therapeutic development. The development of therapeutics that down-regulate a gene is generally easier to achieve than therapeutics that up-regulate a gene; thus, genes with expression levels that are positively correlated with disease risk may be more favorable drug targets for novel therapies. The direction of effect may also provide information to elucidate pathways and the opportunity to explore systems-based approaches to the development of disease. The prediction models can be applied to genotype data of subjects in large biobanks to investigate potential side-effects of drugs with specific gene targets. Finally, direction of effect can be used to improve the interpretation of sequence analyses of genes showing significant correlation of predicted expression with phenotype, since phenotypes associated with reduced expression of genes are more likely to show a relative excess of rare variants. Indeed, we believe that PrediXcan offers intriguing opportunities to combine results of rare and common variant association tests within whole genome sequencing studies, and more generally, to combine results of rare variant gene-based tests from sequencing studies with results of PrediXcan gene-based tests from the large body of existing GWAS for the same phenotypes. Thus, PrediXcan is a method developed to integrate –omics data that can facilitate integration of results from common and rare variant studies.

Regarding the multiple testing correction approach, here we have used Bonferroni correction using the total number of genes tested. In general, both single-variant and PrediXcan analyses will be performed; thus the question that arises is how to address the issue of multiple testing adjustment. The prior probability for a SNP to be causal is much smaller than the prior probability of causality for a gene so it would not be fair to subject SNP tests and gene-based tests to the same level of adjustment. Since we are presenting only gene-based results in our application and given the highly conservative nature of Bonferroni correction, there is no need to further adjust our results. A more conservative approach would be to divide the significance threshold used by a factor of two for the multiple testing using gene-based and SNP-based approaches.

Given the large contribution of regulatory variants on complex traits^9,10,37^, our method is likely to identify causal genes. However, we do not claim causality since SNPs that contribute to the expression of a gene can also act through other mechanisms to determine the phenotype of interest. Replication and experimental validations are needed to determine causality.

In conclusion, we presented a novel gene-based test, PrediXcan that incorporates functional information with regard to gene regulation to identify genes associated with disease traits in large GWAS or whole genome sequence datasets. Our method has the advantage of providing biological insights into the mechanism, namely regulation of gene expression, and direction of effect. This approach can be readily applied to existing GWAS datasets through the use of our publically available PredictDB resource. We further show the utility of our approach by identifying and replicating a number of novel candidate associations within the previously analyzed WTCCC dataset.

## MATERIALS AND METHODS

### Genomic and Transcriptomic Data

#### DGN RNA-Seq Dataset

We obtained whole blood RNA-Seq^38^ and genome-wide genotype data for 922 individuals from the Depression Genes and Networks cohort^16^, all of European ancestry. For our analyses, we used the HCP (hidden covariates with prior) normalized gene-level expression data used for the trans-eQTL analysis in Battle *et al.*^16^ and downloaded from the NIMH repository. Approximately 650K SNPs (minor allele frequency [MAF] > 0.05, Hardy-Weinberg Equilibrium [P > 0.05], non-ambiguous strand [no A/T or C/G SNPs]) comprised the input set of SNPs for imputation, which was performed on the University of Michigan Imputation-Server (https://imputationserver.sph.umich.edu/start.html)^39,40^ with the following parameters: 1000G Phase 1 v3 ShapeIt2 (no singletons) reference panel, SHAPEIT phasing, and EUR population. Non-ambiguous strand SNPs with MAF > 0.05, imputation *R*^*2*^ > 0.8 were retained for subsequent analysis. To reduce computational burden in the application to WTCCC, we used models developed on the HapMap Phase II subset of SNPs.

#### GEUVADIS RNA-Seq Dataset

We obtained freely available RNA-Seq data from 421 lymphoblastoid cell lines (LCLs) generated by the GEUVADIS consortium^15^ and genotype data generated by the 1000 Genomes project (http://www.geuvadis.org/web/geuvadis/RNAseq-project). We used GEUVADIS as a validation dataset to test the gene prediction models generated in the DGN cohort.

#### GTEx RNA-Seq Datasets

We used the nine tissues with the largest sample size in the Genotype-Tissue Expression (GTEx) Pilot Project^14^ to test the gene prediction models generated in the DGN cohort. Tissue samples included subcutaneous adipose (n=115), tibial artery (n=122), left ventricle heart (n=88), lung (n=126), skeletal muscle (n=143), tibial nerve (n=98), skin from the sun-exposed portion of the lower leg (n=114), thyroid (n=112), and whole blood (n=162). In each tissue, normalized gene expression was adjusted for gender, the top 3 principal components (derived from genotype data), and the top 15 PEER factors (to quantify batch effects and experimental confounders)^41^. We used GTEx to test the portability of predictors developed in whole blood (from the DGN cohort) across a wide variety of tissues.

#### Additive model for gene expression traits

We use an additive genetic model to characterize gene expression traits:

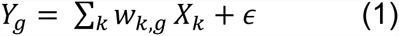

where *Y*_*g*_ is the expression trait of gene *g*, *w*_*k,g*_ is the effect size of marker *k* for gene *g*, *X*_*k*_ is the number of reference alleles of marker *k*, and ε is the contribution of other factors that determine the expression trait assumed to be independent of the genetic component. We note that the summation in model (1) is the genetically determined component of gene expression (i.e., *GRex*).

Effect sizes (*w*_*k,g*_) in model (1) can be estimated using multiple approaches. In this paper we compare penalized approaches such as LASSO (Least Absolute Shrinkage and Selection Operator)^18^ and the elastic net^19^ as well as the more naive simple polygenic score estimates. However, other statistical machine learning approaches^42^, such as Random Forest^43^ or OmicKriging^44^, can be used within the PrediXcan framework to develop prediction models.

The heritability of gene expression defines an upper bound to how well we can predict the trait. We estimated the narrow-sense heritability for each gene using a variance component model with a genetic relationship matrix (GRM) estimated from genotype data, as implemented in GCTA^20^. No pair of subjects from the 922 individuals in DGN shared genetic relatedness 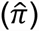 in excess of 5% and thus all were included in the narrow-sense heritability estimation. SNPs in the vicinity of each gene (within 1Mb of gene start or end, as defined by the GENCODE^45^ version 12 gene annotation), with MAF > 0.05, and in Hardy-Weinberg Equilibrium (P > 0.05) were used to construct the GRM for each gene. We calculated the proportion of the variance of gene expression explained by these local SNPs using the following mixed-effects model^37^:

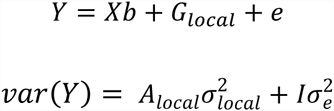

where *Y* is a gene expression trait and *b* a vector of fixed effects. Here *A*_*local*_ is the GRM calculated from the local SNPs, and (the random effect) *G*_*local*_ denotes the genetic effect attributable to the set of local SNPs with var 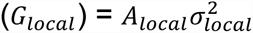. In this paper we focus on the component of heritability driven by SNPs in the vicinity of each gene since the component based on distal SNPs could not be estimated with enough accuracy to make meaningful inferences.

### Estimation of the genetic component of gene expression levels (GReX)

In the simple polygenic score approach, we estimate *w*_*k*_ as the single-variant coefficient derived from regressing the gene expression trait *Y* on variant *X*_*k*_ (as implemented in the eQTL analysis software Matrix eQTL^46^) using the reference transcriptome data. This yields an estimate, 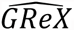, for a GWAS study sample, of the (unobserved) genetically determined expression of each gene *g*:

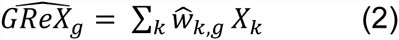

In this implementation of polygenic score, we include all SNPs (regardless of linkage disequilibrium [LD]) that are associated with the expression level of the gene at a chosen p-value threshold in the prediction model.

In contrast, LASSO uses an L1 penalty as a variable selection method to select a sparse set of (uncorrelated) predictors^18^ while the elastic net linearly combines the L1 and L2 penalties of LASSO and ridge regression respectively to perform variable selection^19^. We used the R package *glmnet* to implement LASSO and elastic net with α=0.5 (http://www.jstatsoft.org/v33/i01).

For each gene, LASSO, the elastic net and the simple polygenic score were used to provide an estimate of *GReX* (using equation 2, with the effect size estimates 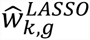, 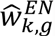, and 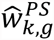, respectively). We included only local SNPs (within 1Mb of the gene start or end). In order to determine the optimal modeling method, we compared the 10-fold cross-validated prediction *R*^*2*^ (the square of the correlation between predicted and observed expression) for the simple polygenic score 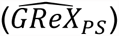 at several p-value thresholds (single top SNP, 1x10^−4^, 0.001, 0.01, 0.05, 0.5, 1) with that from LASSO 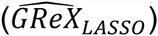 and elastic net 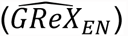.

We also compared the 10-fold cross-validated prediction *R*^*2*^ from elastic net models with different starting SNP sets from the DGN genotype imputation (4.6M 1000 Genomes Project SNPs (MAF>0.05, *R*^*2*^>0.8, non-ambiguous strand), the 1.9M of these SNPs that are also in HapMap Phase II, and the 300K of these SNPs that were genotyped in the WTCCC).

#### Performance of transcriptome prediction in independent cohorts

We tested the feasibility of predicting the transcriptome (i.e., estimating the genetic component of each gene expression trait, 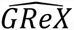, in an independent test transcriptome dataset) using the elastic net effect sizes trained in the DGN whole blood data (n=922). For the test sets, we used independent RNA-Seq datasets from 421 LCL cell lines from the 1000 Genomes project generated by the GEUVADIS consortium^15^ and the nine tissues from the GTEx pilot project^14^ (see Supplemental Figure 3). To assess performance, we used the square of the Pearson correlation, *R*^*2*^, between predicted and observed expression levels.

#### PrediXcan in the WTCCC GWAS Datasets

To illustrate the method, we applied gene prediction models (derived from whole blood) consisting of DGN elastic net predictors to the seven WTCCC disease studies -- bipolar disorder (BD), coronary artery disease (CAD), hypertension (HT), type 1 diabetes (T1D), type 2 diabetes (T2D), Crohn’s disease (CD), and rheumatoid arthritis (RA)^22^. Genotypes imputed to the 1000G reference sets were used. Imputation was done using the University of Michigan Imputation-Server and the same parameters as described for the imputation of DGN data. For each disease, cases and controls (1958 Birth Cohort and the UK Blood Service Cohort) were jointly imputed to avoid subtle differences between cases and controls not attributable to disease risk. We excluded all SNPs with an imputation *R*^*2*^ < 0.8 and for computational speed we kept only the HapMap Phase II subset of SNPs.

For each WTCCC disease, we estimated 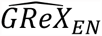, and tested it for association with disease risk using logistic regression in R (R-project.org). We restricted our PrediXcan analysis to include genes with a cross-validated prediction *R*^*2*^ > 0.01 (10% correlation) in the DGN sample. Because the WTCCC studies use shared controls, pleiotropy analyses using these datasets would not be straightforward, and comparison of results across diseases was avoided.

#### GWAS Enrichment analysis

Relative to recent association studies, the WTCCC has a small sample size (∼2,000 cases and ∼3,000 controls per disease). Thus, even with our novel method and a reduced multiple testing burden, our ability to detect numerous novel gene associations may be limited. Alternatively, we tested each disease for an enrichment of known disease genes identified from the NHGRI GWAS Catalog^25^. For each disease, we used the reported genes from the GWAS catalog as the set of known disease genes. We excluded studies listed in the NHGRI GWAS catalog that included the WTCCC samples in order to make sure our known gene lists were independent from the current analysis. We then counted the number of known disease genes that had a PrediXcan p-value below a given threshold. We compared this to the null expectation based on 10,000 randomly drawn gene sets of similar size to the known disease gene set to derive an enrichment p-value. We tested enrichment using PrediXcan p-value thresholds of 0.05 and 0.01.

#### Comparison to large single variant meta-analyses

For the top PrediXcan results in the WTCCC, we cross-referenced the SNPs in the prediction models for these genes with the publically available single-SNP meta-analysis summary results. We excluded T1D from this analysis because, to our knowledge, there are no publically available meta-analysis studies of this disease. We used meta-analyses results for systolic and diastolic blood pressure as a proxy for hypertension. For CD, RA, and BD we were able to use meta-analyses for the same diseases (CD^47^: http://www.ibdgenetics.org/downloads.html, RA^48^, and BD^31^: http://www.med.unc.edu/pgc/downloads).

#### Comparison of gene-based tests (PrediXcan, SKAT, VEGAS)

We compared the results derived from PrediXcan with those from two widely-used gene-based tests, namely VEGAS^49^ and SKAT^50,51^. VEGAS aggregates information from the full set of SNPs within a gene and accounts for LD using simulations from the multivariate normal distribution. SKAT is a kernel-based association test that evaluates the regression coefficients of the SNPs within a gene by a variance component score test in a mixed model framework. We generated BED-formatted files for SNPs and genes (as defined by GENCODE v12) and mapped SNPs that met post-imputation QC parameters to gene regions using bedtools. The use of an offline Perl implementation for VEGAS allowed us to examine the dependence of the results from this approach on LD information through the use of the actual genotype data (versus the default HapMap CEU reference panel data). We developed an R-based pipeline that invokes the SKAT package (version 1.0.1) that is publicly available from CRAN. We generated a Q-Q plot showing the distribution of gene-level p-values for association with RA (for genes outside the HLA region) derived from each gene-based test to test for systematic departure from the null expectation (of uniform p-values).

#### Replication of PrediXcan findings

We selected individuals from Vanderbilt University’s BioVU repository with a diagnosis of rheumatoid arthritis^48^ using a previously validated algorithm for identification of RA cases with a reported positive predictive value of 0.94 and sensitivity of 0.87, as previously described^52^. This trained machine learning classifier was applied to records with at least one International Classification of Diseases, 9th edition code for rheumatoid arthritis to identify true RA cases. RA positive individuals identified by this algorithm were genotyped on two platforms: 833 using the Illumina OmniExpress + Exome chip and 1408 using the Illumina Omni 2.5 BeadChip. A total of 2650 samples from the Illumina Genotype Control set genotyped on Illumina HumanMap550v1/v3 were used for controls. We used the following QC thresholds: sample call rate > 0.98, SNP call rate > 0.99, MAF > 0.05, HWE p-value > 10^−3^. Imputation was performed using IMPUTE2 with the 1000 Genomes phase 1 v3 European samples as the reference panel, phasing was done with SHAPEIT, and SNPs with imputation quality score (“INFO”) > 0.50 were retained. To replicate the PrediXcan RA findings that meet genome-wide significance, we utilized the DGN whole blood elastic net prediction models (as we had done in the discovery WTCCC data). We estimated the genetically regulated gene expression level 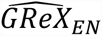 in the replication samples and performed logistic regression with disease status.

## Acknowledgements

We thank Anuar Konkashbaev and Christian Fuchsberger for outstanding technical support and Nicholas Knoblauch for assistance in performing the QC pipeline.

## GTEx data

The Genotype-Tissue Expression (GTEx) Project was supported by the Common Fund of the Office of the Director of the National Institutes of Health (commonfund.nih.gov/GTEx). Additional funds were provided by the NCI, NHGRI, NHLBI, NIDA, NIMH, and NINDS. Donors were enrolled at Biospecimen Source Sites funded by NCI Leidos Biomedical Research, Inc. subcontracts to the National Disease Research Interchange (10XS170), Roswell Park Cancer Institute (10XS171), and Science Care, Inc. (X10S172). The Laboratory, Data Analysis, and Coordinating Center (LDACC) was funded through a contract (HHSN268201000029C) to the The Broad Institute, Inc. Biorepository operations were funded through a Leidos Biomedical Research, Inc. subcontract to Van Andel Research Institute (10ST1035). Additional data repository and project management were provided by Leidos Biomedical Research, Inc.(HHSN261200800001E). The Brain Bank was supported supplements to University of Miami grant DA006227. Statistical Methods development grants were made to the University of Geneva (MH090941 & MH101814), the University of Chicago (MH090951,MH090937, MH101825, & MH101820), the University of North Carolina - Chapel Hill (MH090936), North Carolina State University (MH101819),Harvard University (MH090948), Stanford University (MH101782), Washington University (MH101810), and to the University of Pennsylvania (MH101822). The datasets used for the analyses described in this manuscript were obtained from dbGaP at http://www.ncbi.nlm.nih.gov/gap through dbGaP accession number phs000424.v3.p1.

## WTCCC data

This study makes use of data generated by the Wellcome Trust Case-Control Consortium. A full list of the investigators who contributed to the generation of the data is available from www.wtccc.org.uk. Funding for the project was provided by the Wellcome Trust under award 076113 and 085475.

## DGN data

NIMH Study 7 (GenRED I) - Data and biomaterials were collected in six projects that participated in the National Institute of Mental Health (NIMH) Genetics of Recurrent Early-Onset Depression (GenRED) project. From 1999-2003, the Principal Investigators and Co-Investigators were: New York State Psychiatric Institute, New York, NY, R01 MH060912, Myrna M. Weissman, Ph.D. and James K. Knowles, M.D., Ph.D.; University of Pittsburgh, Pittsburgh, PA, R01 MH060866, George S. Zubenko, M.D., Ph.D. and Wendy N. Zubenko, Ed.D., R.N., C.S.; Johns Hopkins University, Baltimore, MD, R01 MH059552, J. Raymond DePaulo, M.D., Melvin G. McInnis, M.D. and Dean MacKinnon, M.D.; University of Pennsylvania, Philadelphia, PA, RO1 MH61686, Douglas F. Levinson, M.D. (GenRED coordinator), Madeleine M. Gladis, Ph.D., Kathleen Murphy-Eberenz, Ph.D. and Peter Holmans, Ph.D. (University of Wales College of Medicine); University of Iowa, Iowa City, IW, R01 MH059542, Raymond R. Crowe, M.D. and William H. Coryell, M.D.; Rush University Medical Center, Chicago, IL, R01 MH059541-05, William A. Scheftner, M.D., Rush-Presbyterian.

NIMH Study 18 - Data and biomaterials were obtained from the limited access datasets distributed from the NIH-supported “Sequenced Treatment Alternatives to Relieve Depression” (STAR*D). STAR*D focused on non-psychotic major depressive disorder in adults seen in outpatient settings. The primary purpose of this research study was to determine which treatments work best if the first treatment with medication does not produce an acceptable response. The study was supported by NIMH Contract # N01MH90003 to the University of Texas Southwestern Medical Center. The ClinicalTrials.gov identifier is NCT00021528.

NIMH Study 52 (GenRED II) – Data and biomaterials in this release were collected in six projects that participated in the National Institute of Mental Health (NIMH) Genetics of Recurrent Early-Onset Depression (GenRED) project (1999-2009). The Principal Investigators and Co-Investigators were: New York State Psychiatric Institute, New York, NY, R01 MH 060912, Myrna M. Weissman, Ph.D.; Johns Hopkins University, Baltimore, MD, R01 MH059552, J. Raymond DePaulo, M.D., and James B. Potash, M.D., M.P.H.; University of Pennsylvania, Philadelphia, PA (1999-2005), and Stanford University (2006-2009), R01 MH61686, Douglas F. Levinson, M.D. (GenRED coordinator); University of Iowa, Iowa City, IW, R01 MH059542e, Raymond R. Crowe, M.D., and William H. Coryell, M.D.; Rush University Medical Center, Chicago, IL, R01 MH059541-05, William A. Scheftner, M.D.; and University of Pittsburgh, Pittsburgh, PA (1999-2003), R01 MH060866, George S. Zubenko, M.D., Ph.D., and Wendy N. Zubenko, Ed.D., R.N., C.S.

NIMH Study 88 -- Data was provided by Dr. Douglas F. Levinson. We gratefully acknowledge the resources were supported by National Institutes of Health/National Institute of Mental Health grants 5RC2MH089916 (PI: Douglas F. Levinson, M.D.; Co-investigators: Myrna M. Weissman, Ph.D., James B. Potash, M.D., MPH, Daphne Koller, Ph.D., and Alexander E. Urban, Ph.D.) and 3R01MH090941 (Co-investigator: Daphne Koller, Ph.D.).

## Computing resources

This work made use of the Open Science Data Cloud (OSDC) which is an Open Cloud Consortium (OCC)-sponsored project. This work was supported in part by grants from Gordon and Betty Moore Foundation and the National Science Foundation and major contributions from OCC members like the University of Chicago.

https://www.opensciencedatacloud.org/

Grossman RL, Greenway M, Heath AP, Powell R, Suarez R, Wells W, White KP, Atkinson M, Klampanos I, Alvarez H, Harvey C and Mambretti J, The Design of a Community Science Cloud: The Open Science Data Cloud Perspective. (2012) doi:10.1109/SC.Companion.2012.127

This work made use of the Bionimbus Protected Data Cloud (PDC), which is a collaboration between the Open Science Data Cloud (OSDC) and the IGSB (IGSB), the Center for Research Informatics (CRI), the Institute for Translational Medicine (ITM), and the University of Chicago Comprehensive Cancer Center (UCCCC). The Bionimbus PDC is part of the OSDC ecosystem and is funded as a pilot project by the NIH.

https://www.bionimbus-pdc.opensciencedatacloud.org/

Heath AP, Greenway M, Powell R, Spring J, Suarez R, Hanley D, Bandlamudi C, McNerney ME, White KP and Grossman RL, Bionimbus: A Cloud for Managing, Analyzing and Sharing Large Genomics Datasets. J Am Med Inform Assoc (2014) doi:10.1136/amiajnl-2013-002155

---

**Supplemental Figure 1.** Comparison of 10-fold cross-validated predictive performance between all tested methods (LASSO, elastic net with α=0.5, top SNP, polygenic score at several p-value thresholds) in the DGN whole blood cohort. Predictive performance was measured by the *R*^*2*^ between predicted (GReX) and observed expression.

**Supplemental Figure 2.** Comparison of 10-fold cross-validated predictive performance of elastic net in different starting SNP sets (4.6M 1000 Genomes Project (TGP) SNPs, 1.9 M HapMap Phase II SNPs, 300K WTCCC genotyped SNPs) in the DGN whole blood cohort. Predictive performance was measured by the R^2^ between predicted (GReX) and observed expression.

**Supplemental Figure 3. Prediction performance of elastic net in GTEx tissues.** Using whole blood prediction models trained in DGN, we compared predicted levels of expression with observed levels from nine tissues of the GTEx pilot project. The observed squared correlation between predicted and observed gene expression levels, *R2*, is plotted against the null distribution of *R2*.

**Supplemental Figure 4. Comparison of prediction performance between local- and distal-based prediction models.** Using whole blood prediction models trained in DGN, we compared predicted levels of expression with observed levels in GTEx whole blood. Local predictors were generated using elastic net on SNPs within 1Mb of each gene and distal predictors include any *trans*-eQTLs outside this region with a linear regression p<10^−5^. The observed (y-axis) squared correlation between predicted and observed gene expression levels, *R2*, is plotted against the null distribution of *R2*(x-axis).

**Supplemental Figure 5. PrediXcan results in WTCCC.** Q-Q plot of the association p-values from the PrediXcan analysis of 6 remaining WTCCC diseases using expression levels imputed from the DGN whole blood. The red line in each panel shows the null expected distribution of p-values and the blue line represents the bonferroni corrected genome-wide significance threshold. For each disease, the top 3 genes that exceed the bonferroni significance threshold are labeled. The diseases shown are (a) rheumatoid arthritis, (b) Crohn’s disease, (c) bipolar disorder, (d) coronary artery disease, (e) hypertension, and (f) type 2 diabetes.

**Supplemental Figure 6. PrediXcan results in WTCCC.** Plot of the association p-values based on genomic position from the PrediXcan analysis of 6 remaining WTCCC diseases using expression levels imputed from the DGN whole blood. The blue line in each panel represents the bonferroni corrected genome-wide significance threshold. For each disease, the top 3 genes that exceed the bonferroni significance threshold are labeled. The diseases shown are (a) rheumatoid arthritis, (b) Crohn’s disease, (c) bipolar disorder, (d) coronary artery disease, (e) hypertension, and (f) type 2 diabetes.

**Supplemental Figure 7. Enrichment of known disease genes.** Each plot shows the null expected distribution for the number of genes expected to fall below a p-value threshold of 0.01. The null distribution was derived via 10,000 random permutations. The large point on the horizontal axis of each plot shows the observed number of previously known disease genes that fall below the p-value threshold. The diseases shown are (a) rheumatoid arthritis, (b) Crohn’s disease, (c) bipolar disorder, (d) coronary artery disease, (e) hypertension, and (f) type 2 diabetes.

**Supplemental Table 1. Meta-Analysis p-values for SNPs in predictors of top PrediXcan results.** For each of the genes that reached genome-wide significance in our analysis, we looked up the meta-analysis p-values for the SNPs that are included in each of the DNG whole blood predictors. For comparison, we also include the p-value from the single variant analysis of the WTCCC only data.

